# Effects of 7,8-dihydroxycoumarin on the Myelin Morphological Changes and PSD-95 Protein Expression in Balb/c Mice after Sciatic Nerve Injury

**DOI:** 10.1101/529917

**Authors:** Jian Cao, Limin Zhang, Jinlong Li, Hui Leng

**Affiliations:** Department of Orthopedics, Affiliated Hospital of Chifeng University, Chifeng, Inner Mongolia, 024000, China; Department of Ophthalmology, Affiliated Hospital of Chifeng University, Chifeng, Inner Mongolia, 024000, China

**Keywords:** 7,8-dihydroxycoumarin, Peripheral nerve injury, PSD-95 protein

## Abstract

To investigate the effects of 7,8-dihydroxycoumarin on the myelin morphological changes and PSD-95 protein expression in mice with sciatic nerve injury, and to explore the relationship between PSD-95 protein and myelin regeneration after nerve myelin injury. 127 male adult Balb/c mice were selected and randomly divided into high, medium and low 7,8-dihydroxycoumarin dose groups and blank control group. Anastomosis was then carried out for the amputated right sciatic nerve, and intraperitoneal injection of 7,8-dihydroxycoumarin was applied postoperatively. At weeks 1, 2, 4 and 8 after surgery, nervous tissues from the injury side were taken for immunohistochemical Luxol Fast Blue (LFB) staining, so as to observe the morphological changes of the locally injured nerve myelin. Meanwhile, PSD-95 mRNA and protein expression were determined using real-time PCR and western blotting. The nerve myelin recovery in injury side of mice at all time points showed a definite dose-effect relationship with the dose of 7,8-dihydroxycoumarin. Moreover, 7,8-dihydroxycoumarin could inhibit the PSD-95 mRNA level and protein expression. At the same time, there was a dose-effect of the inhibition. 7,8-dihydroxycoumarin can affect nerve recovery in mice with sciatic nerve injury, which shows a definite dose-effect relationship with its dose. Besides, PSD-95 protein expression can suppress the regeneration of the injured nerve myelin.

## 1. Introduction

Peripheral nerve injury is the most common neurological disease in clinic. How to repair and regenerate peripheral nerve injury is a key scientific question in neurosurgery research. At present, clinical peripheral nerve injury mainly includes surgical treatment and drug treatment, but the above two treatment options are not ideal. This is mainly because the nerve tissue itself has the characteristics of high differentiation and low regeneration ability, so the therapeutic effect of peripheral nerve injury repair is still not very ideal. Many studies and clinical trials have shown that natural drugs and extracts can stimulate the expression of nerve growth factor and the proliferation of schwann cells after peripheral nerve injury, thus promoting peripheral nerve regeneration and functional reconstruction[1–4].

7,8-dihydroxycoumarin displays favorable immunosuppression in the immunocompetence screening on natural medicine extracts [5–7]. 7,8-dihydroxycoumarin can dilate coronary blood vessels, increase coronary blood flow, reduce myocardial oxygen consumption, improve myocardial metabolism, promote cardiac function recovery, expand peripheral blood vessels, and prevent arterial thrombosis and inhibit platelet aggregation. At the same time, it has an anti-inflammatory effect on the excitatory pituitary-adrenal cortical system. 7,8-dihydroxycoumarin has regulatory effects on a variety of signaling pathways, including inhibitory protein KB kinase β (IKKβ)/nuclear factors κB (NF-κB) signaling pathway, MAPK signaling pathway, and PI3K/Akt signaling pathway, which play a role in promoting the repair after nerve injury by regulating the above pathways[8–14].

PSD-95 is a postsynaptic density protein, and its discovery has provided a new target for nerve protection treatment after stroke. PSD-95 inhibitor can specifically interfere with the binding of PSD-95 with NMDAR and its downstream signal molecule, block the transduction of excitatory toxicity signal, and provide favorable neuroprotection [15–17]. A large amount of studies had verified that, after peripheral nerve injury, the excessive expression of immune reaction in local injury can partly suppress the nerve repair and regeneration after injury. Typically, the stronger expression of immune reaction would lead to more unsatisfactory nerve fiber regeneration and functional repair. Thus, it can be concluded that, suppressing the excessive expression of immune reaction induced by nerve injury can partially promote the nerve regeneration and repair [18–21].

This paper had detected the PSD-95 protein expression and nerve myelin changes, so as to explore the influence of 7,8-dihydroxycoumarin on the nerve myelin morphological recovery in Balb/c mice after sciatic nerve injury, as well as the correlation between PSD-95 and myelin morphology, thus providing theoretical foundation for the regeneration and repair after peripheral nerve injury.

## 2. Materials and methods

### 2.1 Experimental animals and reagents

A total of 127 8-week-old healthy male Balb/c mice (weight: 16-18 g) were selected from School of Public Health Laboratory Animal Center of Jilin University, among which, 7 died due to anesthesia accident. After unilateral sciatic nerve amputation, the mice were randomly divided into high dose group (H), medium dose group (M), low dose group (L) and blank control group (B), with 30 mice in each group. The blank control group was given 1 mL/d normal saline, while the high, medium and low dose groups were given intraperitoneal injection of 10, 5 and 2.5 mg/kg/d 7,8-dihydroxycoumarin. Finally, 120 mice were enrolled for result analysis.

The major reagents: 7,8-dihydroxycoumarin (with the content of 97.5% identified through HPLC analysis, Jilin Xidian Pharmaceutical Co., Ltd), with the molecular formula of C_9_H_6_O_4_, and the structural formula was shown in Fig.1. Mice PSD-95 western-blot PSD-95 monoclonal antibody (1:1000, Roche Group, Nutley, USA), and Trizol, real-time PCR reagent (Shanghai Sangon Biotechnology Company).

**Fig 1.**
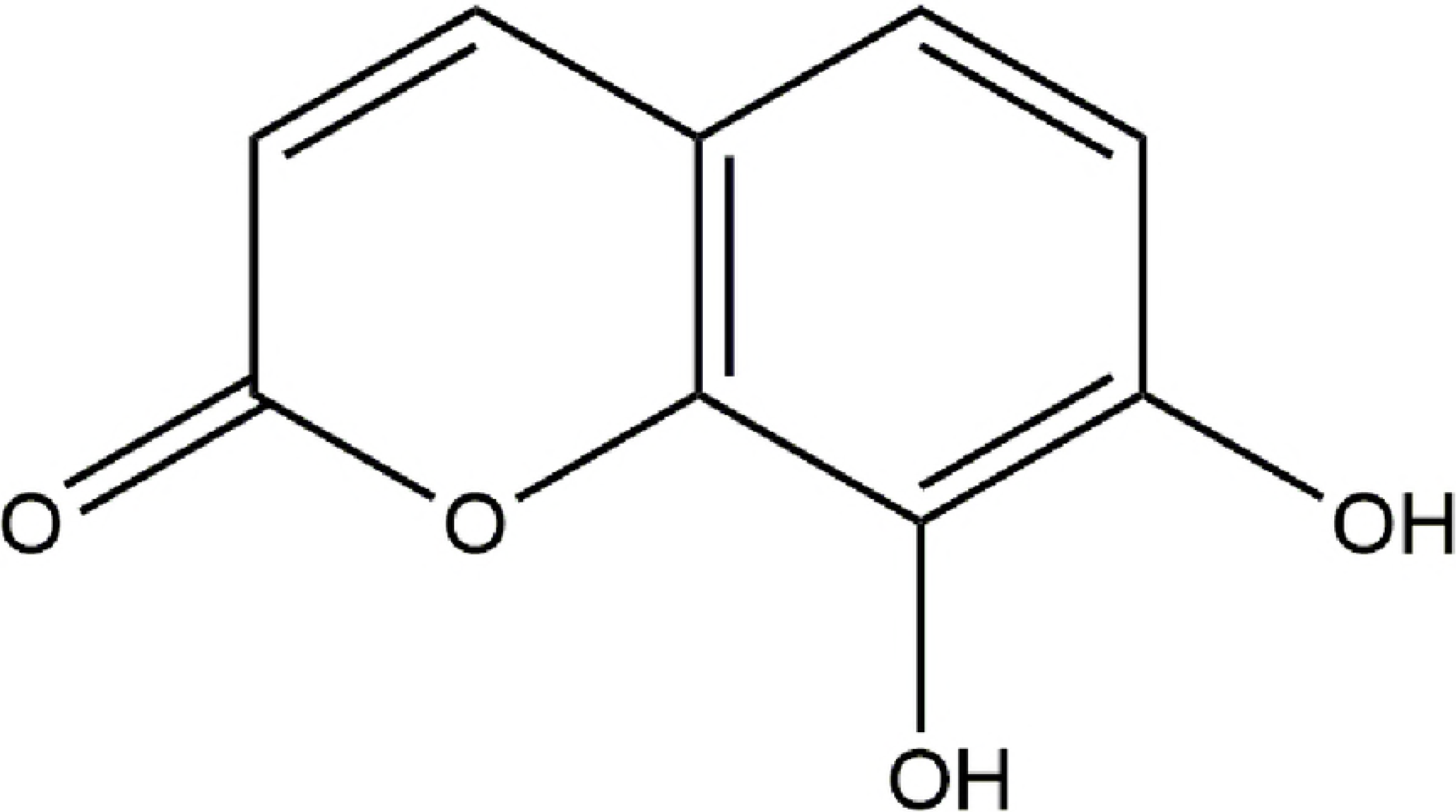
Molecular formula of 7,8-dihydroxycoumarin.

### 2.2 Animal model preparation and administration

The laboratory animals were given intraperitoneal anesthesia using 1% sodium thiopental at the dose of 100 mg/kg. After successful anesthesia, the surgical area was conventionally prepared, an oblique incision was made in the right buttock, and a glass minute hand was used for blunt dissection of muscle, so as to expose the sciatic nerve. The nerve trunk was completely cut off at 0.5 cm below the ischial tuberosity, and the broken ends were aligned under the 12× surgical microscope and sutured using the 11/0 atraumatic stitches. The wound was closed layer-by-layer after washing. Selection of the administration route and dose: intraperitoneal injection was adopted as the administration route. The intraperitoneal injection dose (for 7 consecutive days) of mice was equivalently converted according to the clinical application dose of 7,8-dihydroxycoumarin original medicine. The experimental groups were given intraperitoneal injection of 10, 5 and 2.5 mg/kg/d 7,8-dihydroxycoumarin daily, while the blank control group was injected with normal saline at the same volume. All animal studies were conducted in accordance with the principles and procedures outlined in “Regulations for the Administration of Affairs Concerning Laboratory Animals”, approved by the National Council of China on October 31, 1988, and “The National Regulation of China for Care and Use of Laboratory Animals”, promulgated by the National Science and Technology Commission of China, on November 14, 1988 as Decree No. 2. Protocols were approved by the Committee of Jilin University Institutional Animal Care and Use.

### 2.3 Experimental animal specimen sampling

Five mice were taken from each group at 1, 2, 4 and 8 weeks after modeling for intraperitoneal anesthesia with 1% sodium thiopental at a dose of 100 mg/kg. A posterior median incision was made in the spinal cord, the vertebral plate was opened using the rongeur, and the L_4~6_ spinal cord segments connected with the sciatic nerve were exposed and dissociated intactly. The L_4~6_ spinal cord segments from the injury side was cut, labeled, and rapidly immersed into liquid nitrogen for Western blotting and Real-time PCR analysis. Additionally, sciatic nerve trunk at 0.5 cm from the anastomosis (including the anastomosis) to the distal end was fetched, and fixed in 10% neutral formalin for over 72 h, followed by gradient ethanol dehydration, paraffin embedding and slicing at the thickness of 2 μm.

### 2.4 Real-time PCR assay

The total RNA was extracted using Trizol reagent according the product manual, and the total RNA extracted was used as the template for reverse transcription to prepare the cDNA library. Then, the cDNA library was used as the template to detect the PSD-95 mRNA expression through real-time PCR. The PSD-95 primers were designed using the Beacon designer 9 software (PREMIER Biosoft, USA), with GAPDH as the internal reference, and the primer specificity was verified through Blast (http://blast.ncbi.nlm.nih.gov/Blast.cgi) (Table 1). PSD-95 in tissues from the L_4-6_ spinal cord segments were amplified by PCR (Stratagene M3005p), and a pair of GAPDH primers and Taqman probe were added into each reaction system as the internal reference of PCR amplification. The reaction conditions were as follows: at 95 °C for 30 s, at 58 °C for 60 s, and at 72 °C for 60 s, for a total of 40 cycles. Subsequently, the Ct values of target gene and internal reference gene in each group were detected, which were then incorporated into the standard curve to quantify the target gene, and the histogram was finally drawn (Origin Pro 2017).

**Table 1.**
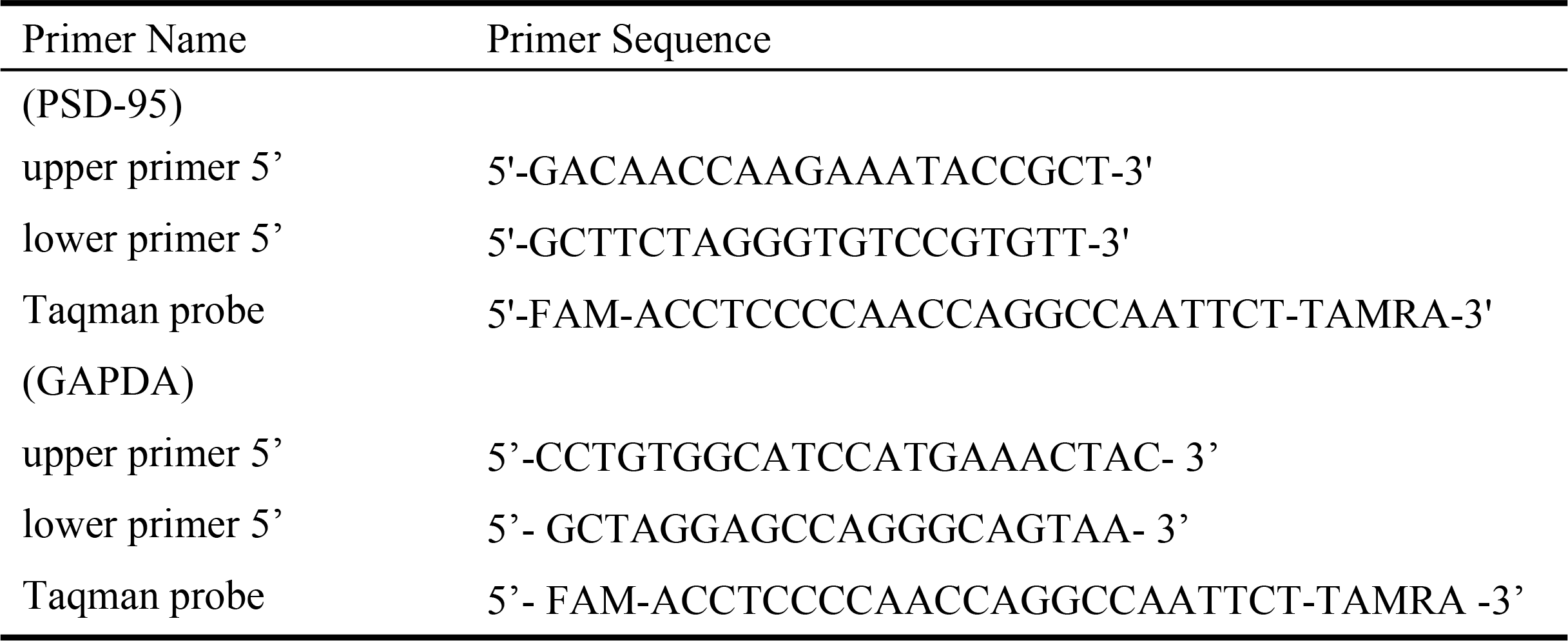
The primers and Taqman probe of PSD-95 and GAPDH.

### 2.5 PSD-95 expression in samples detected through western blotting

Tissues were rapidly taken out from the liquid nitrogen and ground in the mortar; then, RIPA was added to lyse cells, and the proteins were isolated and extracted. Afterwards, the samples were boiled for 15 min in boiling water after adding loading buffer, and the supernatant was taken after centrifugation for 10% SDS-PAGE. The PVDF membranes were treated and immersed into the rabbit anti-mouse PSD-95 monoclonal antibody (1:1000, Roche Group, Nutley, USA) to incubate at 4 °C overnight, followed by washing with 0.01 mol/L PBS for 5 min for 4 times. Later, the samples were immersed into the goat anti-rabbit IgG (1:10000, Roche Group, USA) to incubate at room temperature for 1 h, and washed with 0.01 mol/L PBS for 5 min for 4 times. Color developing was carried out according to the ECL developing kit (Roche Group, USA) instruction, followed by X-ray film exposure. The bands were then scanned and analyzed, the relative molecular weight and absorbance value of the target band were analyzed using the gel image processing system (Alpha-Innotech, California, USA). GAPDF was used as the reference, and a higher absorbance ratio indicated stronger protein expression. Typically, the ratio of absorbance value to internal reference absorbance value was used as the expression index.

### 2.6 Luxol Fast Blue (LFB) staining of myelin

Sciatic nerve trunk at 0.5 cm from the nerve anastomosis (including the anastomosis) to the distal end was taken, and fixed in 10% neutral formalin for over 72 h, followed by gradient ethanol dehydration and paraffin embedding for HE staining. The basic structure of nerve, as well as the changes of with or without cell proliferation and inflammatory cell infiltration were observed, so as to preliminarily screen the objects to be stained. After slicing and deparaffinating, the samples were immersed with LFB staining solution at 60 °C for 12 h, followed by immersion with 95% ethanol for 5 min, addition of 0.05% lithium carbonate for 15 s, washing with 70% ethanol, washing with distilled water, dehydration, transparentizing and sealing. Finally, the samples were observed under the optical microscope (Olympus, Tokyo, Japan), and the number of nerve axon of myelinated nerve fiber were determined using the image scanner (Olympus).

### 2.7 Statistical analysis

The data were presented as the mean values±standard deviations (SD). Statistical analyses were performed using Student’s *t*-test for paired samples and ANOVA for multiple samples. A *p*-value<0.05 was considered statistically significant.

## 3. Results

### 3.1 Real-time quantitative PCR analysis

As shown in Fig.2, changes in the PSD-95 mRNA expression in L_4~6_ spinal cord segments in mice of each groups were detected using real-time PCR. The PSD-95 mRNA contents in corresponding spinal cord segments were elevated after sciatic nerve injury, which showed a gradually increasing trend from the first week to the second week after injury. Moreover, distinct differences could be seen among all groups, among which, the increments in high and medium doses of 7,8-dihydroxycoumarin groups were remarkably lower than those in low dose group and blank control group. 4 weeks later, the PSD-95 mRNA expression shows a decreasing trend in each group. Typically, PSD-95 expression was markedly suppressed in the high and medium dose groups. Taken together, the index detection in each group, changes in PSD-95 mRNA expression in L_4-6_ spinal cord segment tissues in mice at different time points, and myelin LFB staining display a consistent trend. Thus, it can be figured out that, the dose of 7,8-dihydroxycoumarin within the range of 5-10mg/kg/d is beneficial for the repair and regeneration of the injured nerve myelin.

**Fig 2.**
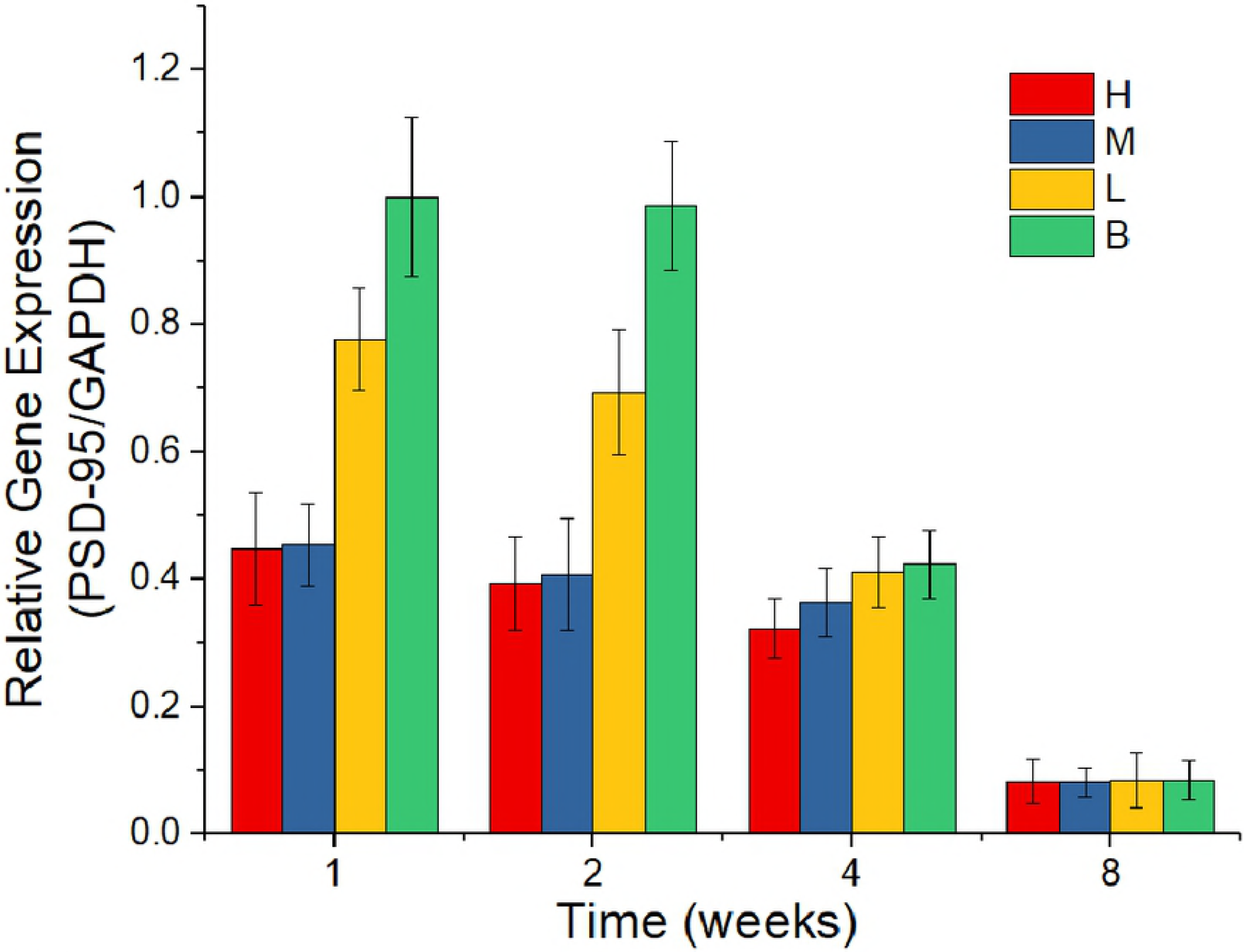
PSD-95 mRNA expression quantities detected by real-time quantitative PCR at 1, 2, 4 and 8 weeks. H (high dose group), M (medium dose group), L (low dose group), B (blank control group).

### 3.2 Western blotting

As shown in Fig.3, changes in PSD-95 protein expression in L_4~ 6_ spinal cord segments in mice of each group were detected through western blotting. At 1 week, the expression of PSD-95 protein in the experiment group was inhibited compared with that in blank control group. The inhibition showed a dose-dependent relationship. The expression level of PSD-95 in groups H, M and L were 70.9%, 23.3% and 17.7% lower than those in group B, respectively. After 2 weeks, the trend was the same as that at 1 week. PSD-95 expression levels in groups H, M and L were reduced by 64.0%, 30.0% and 22.2%, respectively, compared with those in group B. At 4 weeks, the inhibitory rate of 7,8-dihydroxycoumarin on PSD expression levels were 58.5% (H), 27.5% (M) and 15.6% (L), respectively. At week 8, the inhibitory effect was not significant, and the inhibitory rates were 16.7% (H), 9.3% (M) and 10.2% (L), respectively. Comparisons among all groups suggested that, the PSD-95 protein expression was distinctly suppressed in high and medium dose groups at 1-4 weeks. Moreover, compared with the blank control group, 7,8-dihydroxycoumarin at high, medium and low doses could down-regulate PSD-95 protein expression in L_4~6_ spinal cord segments in mice to various degrees.

**Fig 3.**
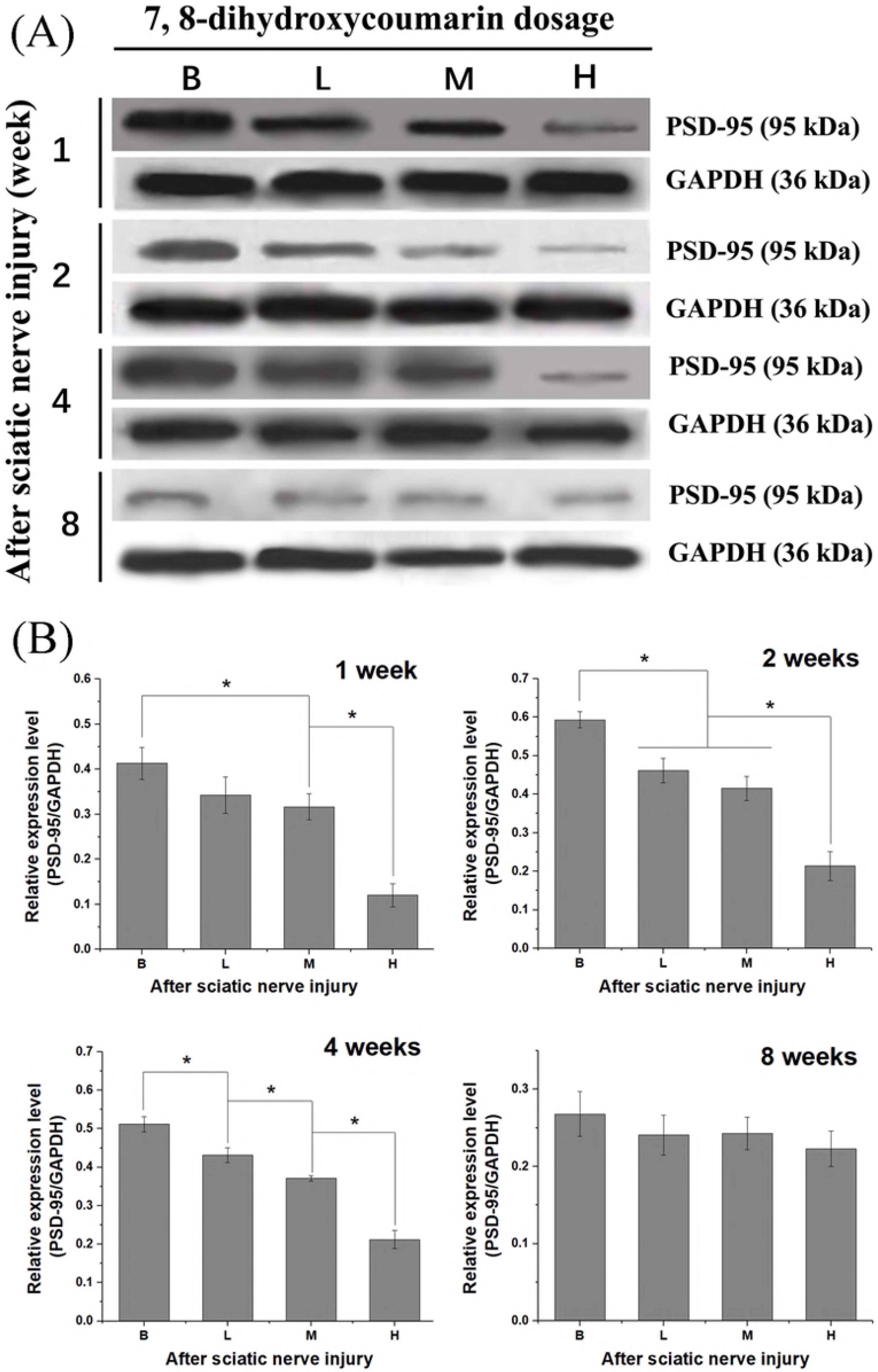
Western blotting was used to quantitatively determine the protein expression of PSD-95 in L_4-6_ spinal cord segments of mice after sciatic nerve injury. (A) Western blotting images, H (high dose group), M (medium dose group), L (low dose group), B (blank control group). (B) According to the gray intensity of bands, the ratios of Psd-95/GAPDH of H, M, L and B groups were calculated, * *P*<0.05, n=3.

### 3.3 LFB staining

LFB staining suggested that, the myelin was azure in color, with the background color of white. 8 weeks after modeling, the myelin in high and medium 7,8-dihydrocoumarin dose groups was regularly arranged, with relatively even myelin thickness, distinct peripheral contour and no obvious fibrous tissue hyperplasia around the myelin. By contrast, the shape and thickness of myelin in low 7,8-dihydrocoumarin dose group were irregular, with clear contour and obvious hyperplasia of nerve tract connective tissues. Moreover, the myelin in blank control group was disorderly arranged, and obvious hyperplastic fibrous and connective tissues could also be observed in Fig.4.

**Fig 4.**
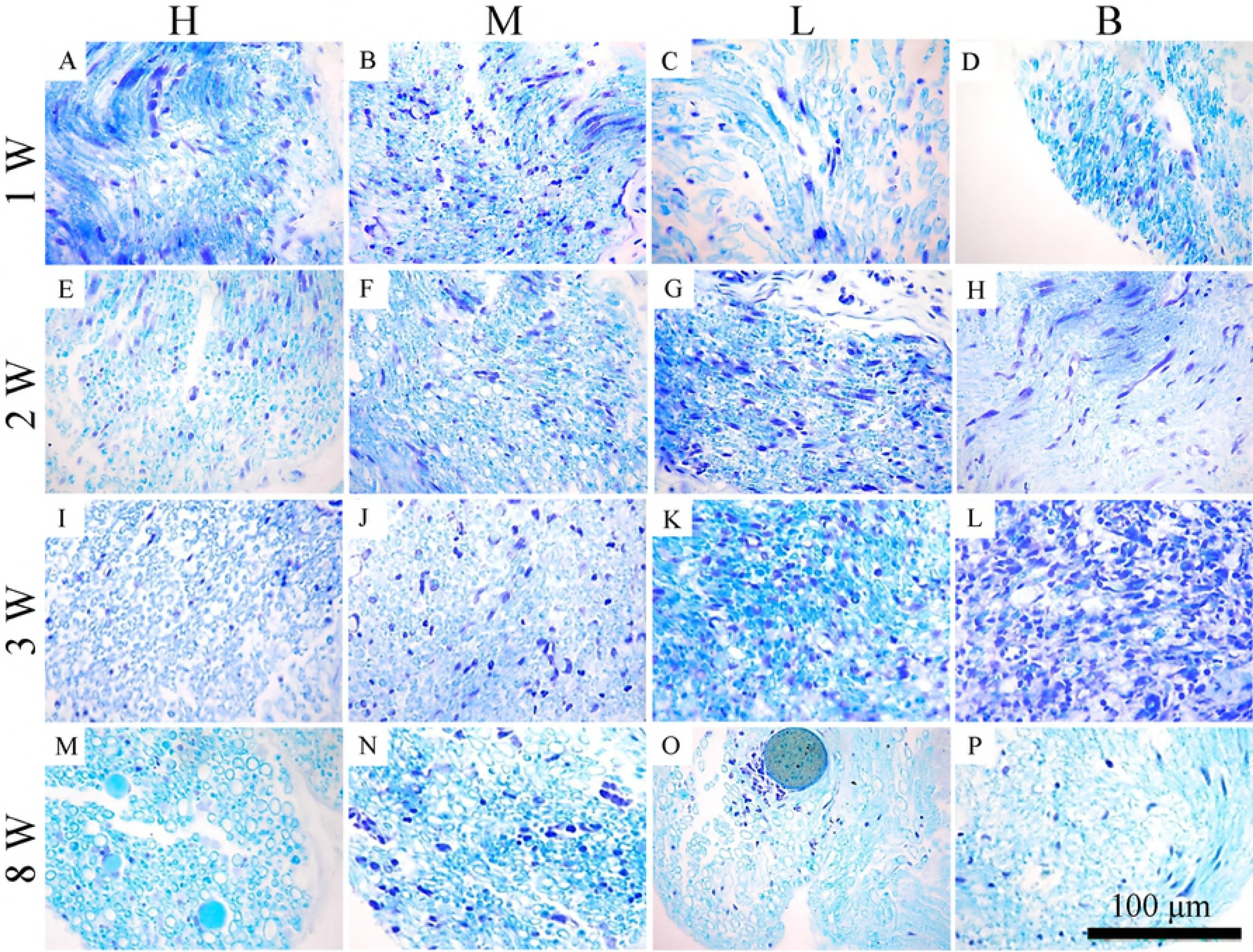
Nerve specimen LFB staining 1 - 8 weeks after modeling. A, E, I and M are high dose group (H); B, F, J and N are medium dose group (M); C, G, K and O are low dose group (L); D, H, L and P are blank control group (B).

Fields of view were randomly selected for image analysis, the number of myelinated nerve fiber were calculated, and the results were statistically analyzed. As shown in Fig.5, there was no significant difference in the number of myelinated nerve fiber between all groups at 1-2 weeks. However, the number of myelinated nerve fiber in the high and medium dose groups were remarkably greater than those in low dose group and blank control group at 4-8 weeks, indicating superior nerve regeneration in high and medium dose groups than in low dose group and blank control group.

**Fig 5.**
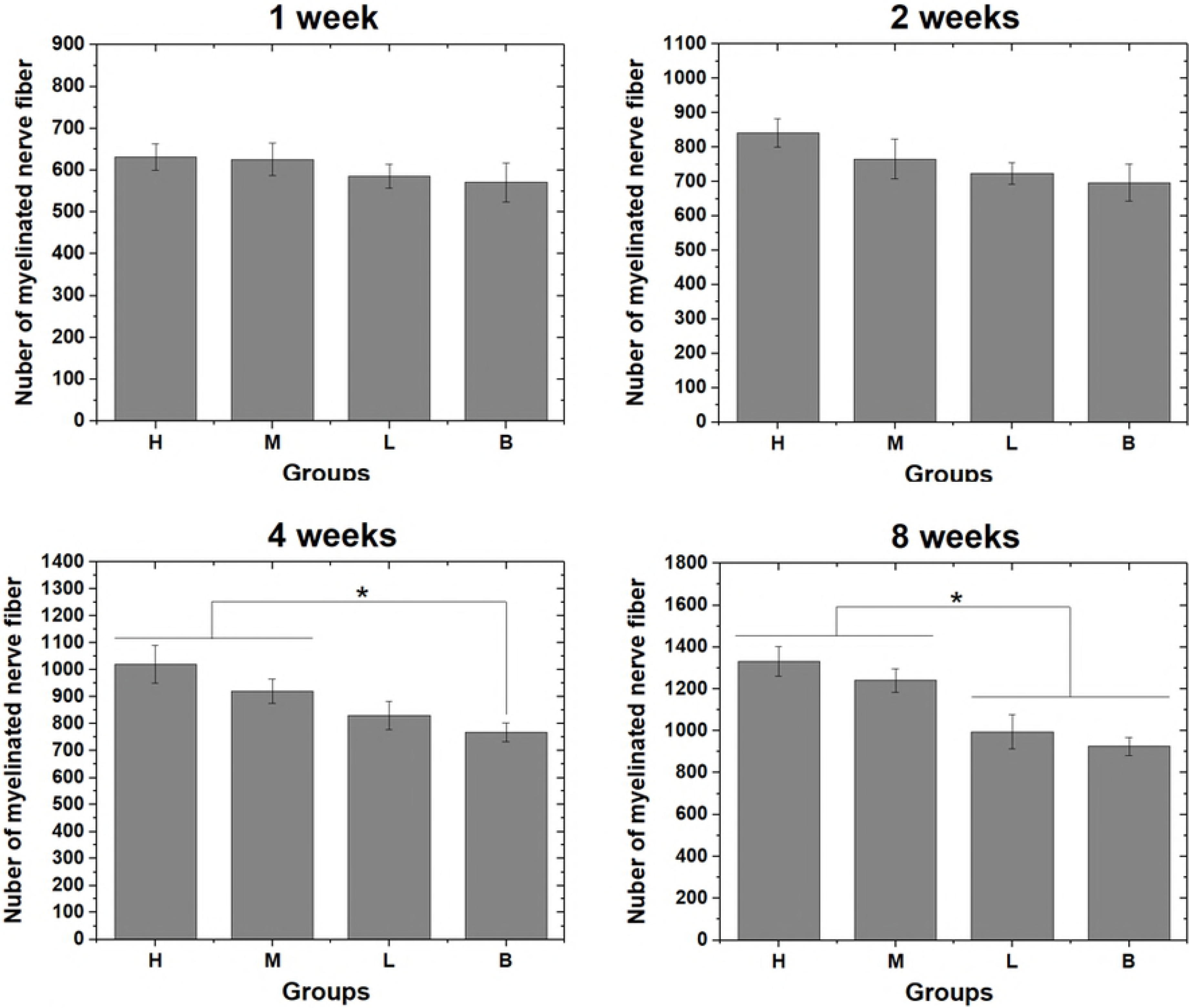
Number of myelinated nerve fiber were calculated in LFB staining images. H (high dose group), M (medium dose group), L (low dose group), B (blank control group) after 1-8 weeks of sciatic nerve injury and 7,8-dihydrocoumarin administration. (*p*<0.05, n=3)

## 4. Discussion

Repair and regeneration after peripheral nerve injury is a long and complicated biological process, which is also one of the challenges to be overcome in surgical field. Drug therapy is an important component of clinical treatment after reconstruction of peripheral nerve continuity. Natural medicine is a kind of multi-element compound, which shows unique advantage to nerve regeneration, since it can potentially provide a more abundant environment with closer proportions of growth active factors to nerve physiological requirements [22, 23]. This paper has treated the 7,8-dihydroxycoumarin as the object of study to observe the influence of 7,8-dihydroxycoumarin on nerve myelin regeneration and postsynaptic density protein-95 (PSD-95) expression in mice after transverse sciatic nerve injury.

PSD-95 is a postsynaptic component protein, which can represent the synaptic morphology and density. Its molecular structure is comprised of 3 repeated PDZ domains in the N-terminal, and the GK domain in the C-terminal, connected by the SH3 domain in the middle. PSD-95 is a structural protein, which shows no protease activity itself, but it can interact with signal molecules (such as NMDA receptor and nNOS) to form a signal complex, thus integrating the excitatory signal at synaptic level. Moreover, PSD-95 can interact with neurexins (presynaptic membrane overhanging protein) through neuroligins (the postsynaptic neuron adhesion molecule), thus participating in connecting the synaptic structure and maintaining its function. Research verifies that, PSD-95 can render the more stable interacting membrane proteins at synaptic level; meanwhile, it can regulate the functional characteristics of these membrane proteins, which manifests that the NMDA receptor gated pathway shows PSD-95 dose-dependent changes as well as the inward integration of PSD-95 on the potassium channel[24]. Tao et al. discovered in their study that, a large amount of PSD-95 expression was found in the neuron apoptosis animal model induced by ischemic cerebral injury and excitatory toxicity[25]. However, the excess NO production resulted from the enhanced nNOS activity is one of the major factors leading to post-ischemic nerve toxic injury [26]. Suppressing PSD-95 expression or blocking the binding of PSD-95 with NMDA receptor shows protection for excitatory toxicity and ischemic brain injury [15]. PSD-95 can exert injury to peripheral nerve through the NMDA receptor-NO pathway [27].

In this study, mice are given intraperitoneal injection of 7,8-dihydroxycoumarin after sciatic nerve injury and are monitored for as long as 8 weeks. Real-time PCR and Western blotting results indicate that, PSD-95 expression is suppressed in the spinal cord segments of high and medium 7,8-dihydroxycoumarin dose groups, which is remarkably lower than that in low dose group and blank control group. Besides, results of LFB staining on nerve myelin further demonstrate that, the regeneration of injured nerve is accompanying with the increases in the number, integrity and thickness of nerve myelin. As shown in the results, the number and average diameter of myelinated nerve fiber in high and medium dose groups are markedly lower than those in low dose group and blank control group; besides, those in low dose group are also notably higher than those in blank control group. These findings suggest that 7,8-dihydroxycoumarin show protective effect on the myelin growth of the injured nerve. The low dose group can not effectively restrain PSD-95 protein expression (no significant difference compared with the blank control group), but it also exerts obvious protection on the myelin growth of the injured nerve, and such protection mechanism is associated with the suppression of PSD-95 protein expression.

A phenomenon can also be observed in this study, which is that, after continuous injection of 7,8-dihydroxycoumarin for 8 weeks, no significant difference in PSD-95 expression can be seen when comparing the medium and high dose groups with the low dose group and blank control group, while significant difference can be observed in nerve myelin status. Such finding may reveal that, the nerve repair and regeneration process has been completed in medium and high dose groups after 8 weeks. Subsequently, the PSD-95 expression level is decreased, which approaches that in low dose group and blank control group, demonstrating the completion of nerve repair and regeneration 8 weeks after sciatic nerve injury.

With regard to the mechanism by which 7,8-dihydroxycoumarin promotes nerve repair after injury, this study finds that, 7,8-dihydroxycoumarin application after peripheral nerve injury in mice can suppress PSD-95 mRNA expression. Besides, the PSD-95 expression quantity is progressively decreased with the increase in 7,8-dihydroxycoumarin dose. On the other hand, LFB staining results in local injury suggest that 7,8-dihydroxycoumarin application leads to markedly superior local nerve myelin recovery to that in blank control group. Thus, we speculate that, 7,8-dihydroxycoumarin may alleviate the demyelination degree during nerve repair and regeneration through suppressing the continuous PSD-95 expression after nerve injury and reducing the excess NO production-induced neural toxic injury, which has thereby provided favorable conditions for nerve repair and regeneration after injury.

This study is associated with certain drawbacks. Typically, correlation analysis is only performed between the morphological changes of nerve myelin after peripheral nerve injury and the changes in PSD-95 protein expression, but the local immune reaction after nerve injury is not detected. Therefore, future study should further intensively investigate the expression of post-injury nerve regeneration-related proteins (p75NTR and S100β), inflammatory reaction-associated cellular signaling pathways, and the mechanism of action by which 7,8-dihydroxycoumarin mitigates the neurotoxic injury resulted from excess NO production after nerve injury.

## 5. Conclusions

In this study, the sciatic nerve injury model of Balb/c mice was used to study the effects of 7,8-dihydroxycoumarin on peripheral nerve injury regeneration. Through to the sciatic nerve injury in mice given different concentrations of 7, 8-dihydroxycoumarin, realtime PCR and western-blot results indicated that PSD - 95 mRNA and expression level could be significantly suppressed at 1-2 weeks timepoint. The inhibitory effect of 7, 8-dihydroxycoumarin on PSD-95 was dose dependent. The concentration of 5-10 mg/kg/d 7,8-dihydroxycoumarin inhibited PSD-95 expression most significantly. Inhibition of PSD-95 expression after nerve injury can specifically interfere with the binding of PSD-95 to NMDAR and its downstream signal molecules, block the conduction of excitatory toxicity signal, and provide good neuroprotective effect. Therefore, it was determined that 7,8-dihydroxycoumarin could promote nerve repair and regeneration of myelin in injured sciatic nerve in mice.

## Author contributions

JC and Lm Z designed the study. Jl L performed western blot. HL carried out the animals experiments. Lm Z analyzed the data. JC wrote the paper.

